# Microglial phagocytosis mediates long-term restructuring of spinal GABAergic circuits following early life injury

**DOI:** 10.1101/2023.01.29.525735

**Authors:** Yajing Xu, Dale Moulding, Wenanlan Jin, Simon Beggs

**Affiliations:** University College London; UCL GOS Institute of Child Health

## Abstract

Peripheral injury during the early postnatal period alters the somatosensory system, leading to behavioural hyperalgesia upon re-injury in adulthood. Spinal microglia have been implicated as the cellular mediators of this phenomenon, but the mechanism is unclear. We hypothesised that neonatal injury (1) alters microglial phagocytosis of synapses in the dorsal horn leading to long-term structural changes in neurons, and/or (2) trains microglia, leading to a stronger microglial response after re-injury in adulthood. Using hindpaw surgical incision as a model we showed that microglial density and phagocytosis increased in the dorsal horn region innervated by the hindpaw. Dorsal horn microglia increased engulfment of synapses following injury, with a preference for those expressing the vesicular GABA transporter VGAT and primary afferent A-fibre terminals in neonates. This led to a long-term reduction of VGAT density in the dorsal horn and reduced microglial phagocytosis of VGLUT2 terminals. We also saw an increase in apoptosis following neonatal injury, which was not limited to the dorsal horn suggesting that larger circuit wide changes are happening.

In adults, hindpaw incision increased microglial engulfment of predominantly VGAT synapses but did not alter the engulfment of A-fibres. This engulfment was not affected by prior neonatal injury, suggesting that microglial phagocytosis was not trained. These results highlight microglial phagocytosis in the dorsal horn as an important physiological response towards peripheral injury with potential long-term consequences and reveals differences in microglial responses between neonates and adults.

## Introduction

The neonatal period is a critical period during which perturbations in neural activity can cause long-lasting changes (Hensch, 2005). For the somatosensory system, both clinical and animal studies have shown that tissue injury during the early postnatal period leads to long-term alteration in somatosensory processing (Schwaller & Fitzgerald, 2014). For example, subjects with a history of neonatal injury exhibit hyperalgesia upon re-injury in adulthood (Peters et al., 2005; Ren et al., 2004; Taddio et al., 1997; Walker et al., 2009), but only if the first injury happened during a critical period in postnatal development (Walker et al., 2009).

The neuronal underpinnings of these acute and long-term behavioural changes following injury have been studied in animal models following peripheral hindpaw incision injury (a model of post-operative pain) (Brennan et al., 2005; Walker et al., 2016), but what governs the neuronal activity changes in this process is still unclear. We have previously shown that spinal microglial inhibition prevents neonatal-incision induced hyperalgesia in adulthood, suggesting that microglia play a causative role in sensitising dorsal horn circuits long-term (Beggs et al., 2012; Moriarty et al., 2019).

Microglia regulate developmental pruning of synapses in the brain and spinal cord (Paolicelli et al., 2011; Schafer et al., 2012a; Vainchtein et al., 2018; Vukojicic et al., 2019), and we have recently shown that microglia are critical for the postnatal refinement of sensory A-fibre projections in the dorsal horn (Xu et al., 2021). In addition, it has been shown that microglia cells can exhibit innate immune memory, by which a stimulus causes microglia to enter a ‘trained’ state which alters their future response to inflammatory challenges (Neher & Cunningham, 2019).

Therefore, we hypothesised that neonatal injury might (1) alter microglial pruning of synapses acutely, leading to long-term neuronal changes and/or (2) ‘train’ microglia and alter their response long-term towards subsequent injury in adulthood. Both possibilities (long-term neuronal changes vs. long-term microglial changes) could underly the long-term functional and behavioural changes observed following neonatal injury.

To address these questions, we used hindpaw incision as a model of neonatal surgical injury and quantified microglial engulfment of local excitatory and inhibitory synapses and primary afferent A-fibre central axonal projection in the dorsal horn following incision in neonates, and in adults with or without prior neonatal incision. Neonatal incision increased apoptosis in the spinal cord, and acutely increased microglial engulfment of inhibitory VGAT presynaptic terminals and A-fibres following incision, but not excitatory VGLUT2 presynaptic terminals.

This resulted in a long-term decrease of inhibitory synapse density in adulthood, and a reduction of VGLUT2 engulfment by microglia. In contrast, adult incision acutely increased microglial engulfment of predominantly inhibitory presynaptic terminals and to a lesser degree excitatory presynaptic terminals, but not A-fibres. The injury induced phagocytosis in adults was not altered by neonatal incision, suggesting that microglial phagocytosis was not trained.

## Materials and Methods

### Animals

Sprague-Dawley rats and transgenic mice on C57BL/6J background of both sexes were used. For visualisation of A-fibres, Slc17a7-IRES2-Cre (Vglut1-Cre) males (Jackson Laboratory 023527) were crossed with Ai9 females (Jackson Laboratory 007909) to obtain animals that expressed the tdTomato fluorophore under the Vglut1 promoter (*Vglut1^Cre/+^*; *R26^LSL-Ai9/+^*) (Chamessian et al., 2019; Todd et al., 2003; Yasaka et al., 2014).

Data points are presented as black (female) or red (male) to indicate the sexes. Numbers of animals used were based on previous experiments and are indicated in the figures for each experiment. For a table with detailed species, ages, sexes, and numbers of animals used in each experiment, please see Supplementary Table 1. All procedures were carried out in accordance with the guidelines of the UK Animals (Scientific Procedures) Act 1986 and subsequent amendments.

### Hindpaw incision injury

P3 rats or P4-P5 mice were induced with 5% isofluorane anaesthesia and maintained at 3.5% isoflurane through a nose-cone. Body temperature of the animal was maintained with a heating pad throughout the procedure. The plantar skin of the left hind paw was incised from midpoint of the heel to the first foot pad, and the flexor digitorum brevis muscle underneath was lifted and transected. The wound was then closed with surgical glue, and animals were allowed to make full recovery from the anaesthesia before returning to their cages. Control animals received only anaesthesia without hindpaw incision to control for any effects of anaesthesia. This is a modification of the well-established model of surgical pain (Brennan et al., 1996).

### Immunohistochemistry

Animals were overdosed with pentobarbital and transcardially perfused with saline followed by ice-cold 10% formalin. The sciatic nerve was exposed and traced to locate L4 & L5 dorsal root ganglia (DRG) and the corresponding region of the lumbar spinal cord was dissected and post-fixed in 10% formalin overnight, followed by immersion in 30% sucrose until they sank. 50μm free-floating spinal cord sections were cut on the microtome with every 2nd section collected.

Tissue sections were washed 3 × 10 min in PBS and then incubated in blocking solution (10% donkey serum, 0.2% Triton X-100 in PBS) for 2.5h at room temperature. The sections were then incubated with primary antibodies at 4°C overnight followed by secondary antibodies at room temperature for 2h, both diluted in blocking solution (3% donkey serum, 0.2% Triton X-100 in PBS) (for list of antibodies and their respective concentrations used, see Supplementary Table 2). Samples were mounted in Fluoromount Aqueous Mounting Medium (Sigma) or ProLongTM Diamond Antifade Mountant (Thermo Fischer), if the tissue contained endogenous fluorophores.

### Image acquisition and analysis

Confocal z-stacks were taken with a Zeiss LSM880 confocal microscope using a 20× water immersion objective (NA 1.0, pixel size 0.3 μm (x) × 0.3 μm (y) × 0.67 μm (z)) for imaging of A-fibres and 63× oil immersion objective (NA 1.4, Nyquist resolution pixel size 0.07 μm (x) × 0.07 μm (y) × 0.23 μm (z)) for imaging of synapses. For neonates, the full thickness of the section was imaged. In adults, the VGAT and VGLUT stain did not fully penetrate the sections, so only the top 5um from the imaging surface were acquired. Images were cropped as needed to a 192 × 192 μm field of view to include only the superficial dorsal horn laminae for analysis. Images taken with the 63x objective were additionally deconvolved using the Huygens software. Apoptotic cell numbers were manually counted using an epifluorescent wide-field microscope (Leica DMR, 20x air objective, NA 0.5) with the experimenter blinded to experimental conditions. All image acquisition details can also be found in Supplementary Table 1.

Only intact sections with an even stain were analysed, and at least 6 sections were imaged and analysed per animal to reduce variability. A-fibre and synapse engulfment by microglia was analysed with automated batch processing in Fiji and the 3D-ROI manager plugin (Linkert et al., 2010; Ollion et al., 2013; Schindelin et al., 2012; Schneider et al., 2012). This procedure binarised each of the channels containing staining for microglia/lysosomes/A-fibres or synapses and measured the volume of their overlap (individual macros were written for A-fibre, VGLUT2 and VGAT analysis respectively). Raw data and scripts for the automated analysis are available online at https://www.ebi.ac.uk/biostudies/ under the accession number S-BSST1009 (https://www.ebi.ac.uk/biostudies/studies/S-BSST1009).

### Experimental design

#### Neonatal incision (Fig. 1a)

Neonatal hindpaw incision was performed on Sprague Dawley rats at P3 and one day later on mice at P4-P5 to facilitate handling, as mouse pups are much smaller than rat pups and still very delicate at P3. The lumbar spinal cord was extracted 3 days later at P6 or P7 for rats and mice respectively, as that has been shown to be the time of maximal microglial proliferation in adults (Beggs et al., 2012; Obata et al., 2006; Wen et al., 2009). Age matched control litters received only anaesthesia without incision. Each experimental group consists of animals pooled from at least two different litters to avoid litter effects.

**Figure 1.**
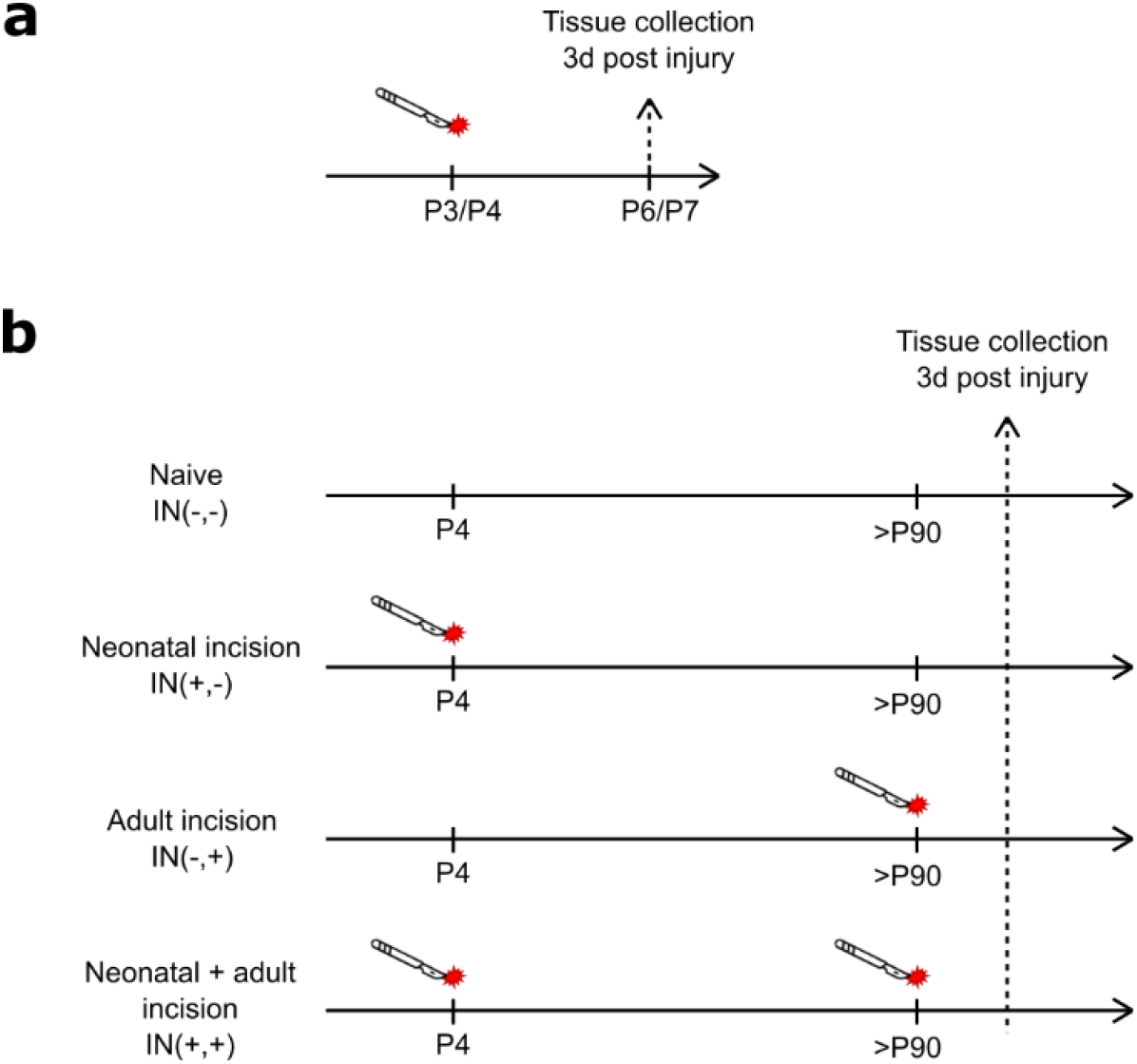
Experimental design for neonatal (a) and adult (b) incision. The lumbar cord (L4-L5) was extracted three days after adult incision in IN(−,+) and IN(+,+) animals or at equivalent age in animals without adult incision in IN(−,−) and IN(+,−). All analysis were carried out in the medial superficial dorsal horn that is innervated by the hindpaw. Males and females were pooled together for analysis to increase the statistical power, but data points are presented as black (female) or red (male) to indicate the sexes.

#### Adult incision (Fig. 1b)

Unless stated otherwise, 4 experimental groups of mice were used in all experiments, with different combination of neonatal and adult incision treatment:

- Naive (IN(−,−)) animals did not receive any treatment
- Neonatal incision only animals (IN(+,−)) received neonatal anaesthesia and incision, but no adult anaesthesia or incision
- Adult incision only animals (IN(−,+)) received neonatal anaesthesia and adult incision under anaesthesia
- Neonatal and adult incision animals (IN(+,+)) received both neonatal incision and adult incision both under anaesthesia.

### Statistical Analysis

Null-hypothesis significance testing were carried out in GraphPad Prism 6. Two-way ANOVA was used for comparisons across two factors, One-way ANOVA was used in Fig. S1 to compare different conditions. Welch’s t-test was used in Fig. 4. Where applicable, post-hoc comparisons were made using the Sidak method. Significance level was set at α = 0.05. N-numbers and P-values are indicated in text and figures.

Estimation statistics for the 95% confidence intervals (95% CI) of the mean difference were calculated on estimationstats.com (Ho et al., 2019) using 5000 samples of bias-corrected and accelerated bootstrapping. Additionally, conventional null-hypothesis significance testing were carried out on estimationstats.com and GraphPad Prism 6 for all comparisons (significance level was set at α=0.05), which are listed in the Supplementary Table 1.

Data are presented as mean ± SD in all figures. The effect size is presented as 95% CI of the mean difference on a separate but aligned axis. The mean difference is plotted as a dot on the background of its probability distribution, and the 95% confidence interval is indicated by the ends of the error bar. All values in text and figures are given with two decimals or rounded to two significant figures. For a comprehensive list with exact statistical values and analyses, see Supplementary Table 1.

## Results

### Neonatal incision acutely increases microglial engulfment of inhibitory, but not excitatory presynaptic terminals

The medial L4-L5 dorsal horn ipsilateral to the incision site was analysed for microglial changes in cell and lysosome volume as measured by Iba1 and CD68. Following incision, a significant increase in microglial volume was observed for both sexes, suggesting a proliferation of microglia cells (F (1, 19) = 5.45, P = 0.031) (Fig. 2a, b). No lysosomal volume change was detected following incision (Fig. 2a, c) (F(1,20) = 0.71, P = 0.41). Females had higher volumes of microglia and lysosomes compared to males (F(1,19) = 5.03, P = 0.037 and F(1,20) = 6.21 P = 0.022), but there was no statistical interaction between sex and incision (F(1,19) = 0.18, P = 0.67 and F (1, 20) = 1.40, P = 0.2502).

**Figure 2.**
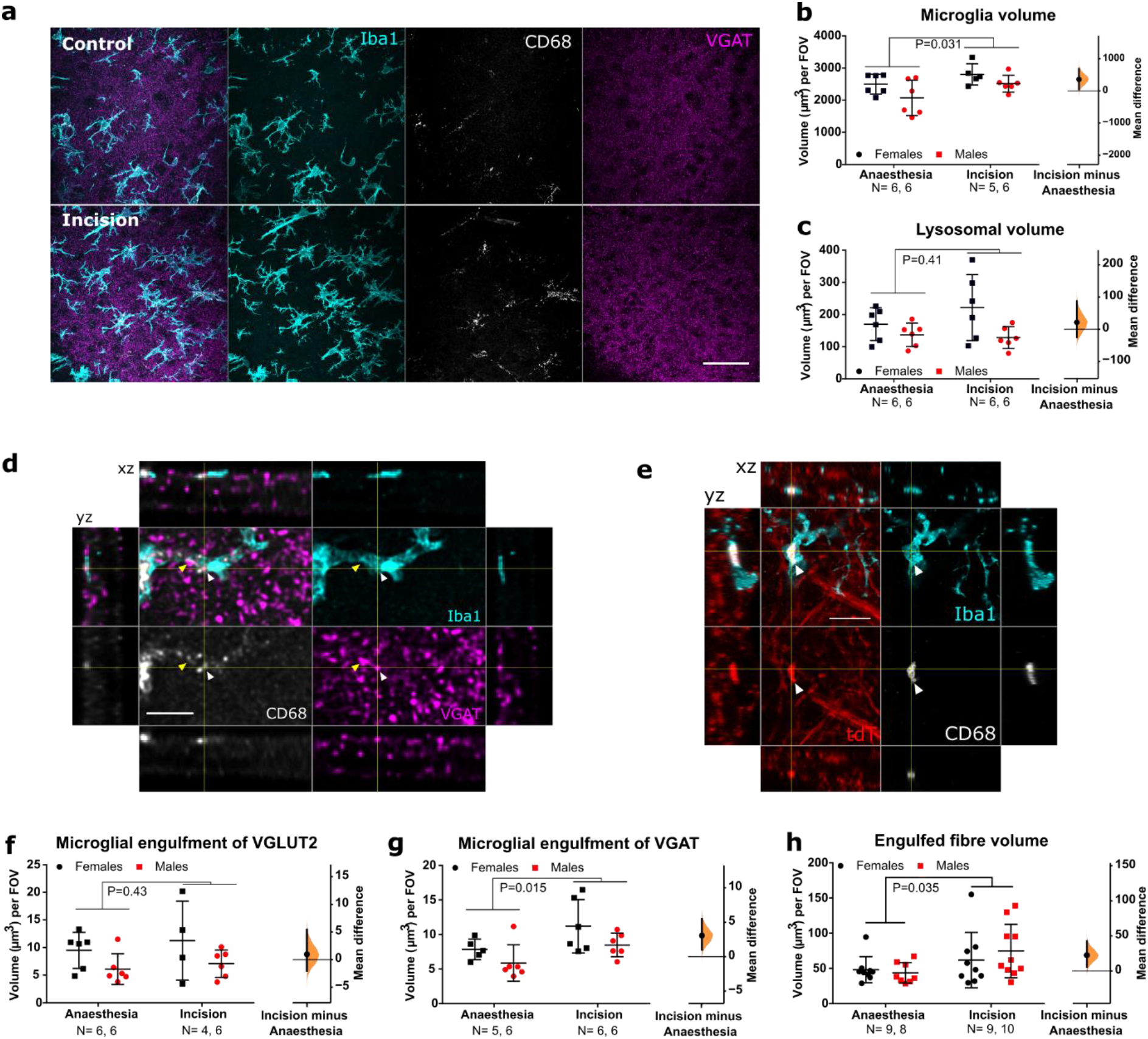
Neonatal incision acutely increases microglial engulfment of VGAT and A-fibres. **a**.Representative maximum projection of confocal images from the dorsal horn showing microglia (Iba1, cyan), lysosomes (CD68, grey), and inhibitory synapses (VGAT, magenta) from a control (top panels) and incision animal (bottom panels). Scale bar = 50μm. **b**. Microglial volume increases following incision, treatment mean difference 359.08 [95.00% CI 49.45, 690.24]. **c**. Lysosomal volume within microglia did not increase significantly, treatment mean difference 21.40 [95.00% CI −25.15, 87.34]. **d**. Representative confocal images from the dorsal horn with arrow heads showing co-localisation of inhibitory synapses (VGAT, magenta) with lysosomes (CD68, grey) inside microglia (Iba1, cyan). Cross-hairs show position of the xz and yz side-view panels. Scale bar = 5μm. **e**. Representative confocal images from the dorsal horn with arrow heads showing co-localisation of A-fibres (tdT, red) with lysosomes (CD68, grey) inside microglia (Iba1, cyan). Cross-hairs show position of the xz and yz side-view panels. Scale bar = 5μm. **f**. Neonatal incision did not significantly affect microglial engulfment of VGLUT2 synapses. Treatment mean difference 0.95 [95.00% CI −2.02, 5.38] **g**. Neonatal incision significantly increased the engulfment of VGAT synapses. Treatment mean difference 3.07 [95.00% CI 1.04, 5.48]. **h**. Neonatal incision significantly increased the engulfment of A-fibres. Treatment mean difference 22.56 [95.00% CI 5.78, 42.30]. N-numbers and P-values for two-way ANOVA are indicated in figures. Field of view (FOV) = 192.79 × 192.79 × 50 μm

Next, we asked whether neonatal incision acutely alters microglial interaction with local excitatory and inhibitory synapses of interneurons in the dorsal horn. As microglia are known to primarily engulf presynaptic terminals (Weinhard et al., 2018), we examined microglial engulfment of excitatory (VGLUT2) and inhibitory (VGAT) presynaptic markers of spinal interneurons (Punnakkal et al., 2014). There was no change in the microglial engulfment of excitatory VGLUT2 (F(1,18) = 0.65, P = 0.43), but an increase in the engulfment of inhibitory VGAT was observed for both sexes following incision (F (1, 19) = 7.25, P = 0.014) (Fig. 2d, f, g). Females had higher levels of engulfment for both VGLUT2 and VGAT than males (F(1,18) = 4.95, P = 0.039 and F(1,19) = 4.46, P = 0.048 respectively), but there was no interaction between sex and treatment (F (1, 18) = 0.051, P = 0.82 and F (1, 19) = 0.11, P = 0.74 respectively).

### Neonatal hindpaw incision acutely increases microglial engulfment of A-fibre central projections

Having established that microglia selectively engulf inhibitory presynaptic terminals of local interneurons, we asked whether injury also alters engulfment of afferent A-fibres in the dorsal horn, given our previous finding that microglia prune A-fibres during this period (Xu et al., 2021). To visualise A-fibres, *Vglut1^Cre/+^*∷ *R26^LSL-Ai9/+^* mice were used which express the tdTomato (tdT) fluorophore under the *Vglut1* promoter, and labels a subset of A-fibres, which are myelinated low-threshold mechanosensitive afferents (Chamessian et al., 2019; Todd et al., 2003; Yasaka et al., 2014). Significantly more tdT labelled A-fibres were engulfed by microglia after incision compared to anaesthesia controls for both sexes (F (1, 32) = 4.85, P = 0.035) (Fig. 2 e, h). There was no significant sex effect (F (1, 32) = 0.17, P = 0.69) or interaction between sex and treatment (F (1, 32) = 0.75, P = 0.39).

### Neonatal hindpaw incision increases spinal apoptosis

In addition to microglial interaction with synapses we also investigated the effect of neonatal incision on apoptosis. Neonatal incision is known to increase apoptosis in the spinal cord (Moriarty et al., 2019), and as microglia are capable of inducing apoptosis both during development and in adulthood (Marín-Teva et al., 2011), we hypothesised that apoptotic cells might accumulate in the medial dorsal horn, which received afferent input form the hindpaw and where microglia cells become reactive following incision. To investigate this, we counted and mapped the location of caspase-3 labelled apoptotic cells in the ipsilateral hemisections from the lumbar region and normalised to the number of sections counted. Neonatal incision significantly increased the number of apoptotic cells in the ipsilateral L4-L5 spinal cord in both sexes (F (1, 18) = 6.32, P = 0.022) compared to anaesthesia controls (Fig. 3b). The location of apoptotic cells was not just in the dorsal horn but highest in three distinct regions: the superficial lateral dorsal horn, along the central canal, and in the ventral horn (Fig. 3a). This suggests that incision-induced apoptosis is likely unrelated to direct incision-induced microglial reactivity in the dorsal horn. There was no interaction or sex effect (F (1, 18) = 0.28, P = 0.60 and F (1, 18) = 0.092, P = 0.77 respectively).

**Figure 3.**
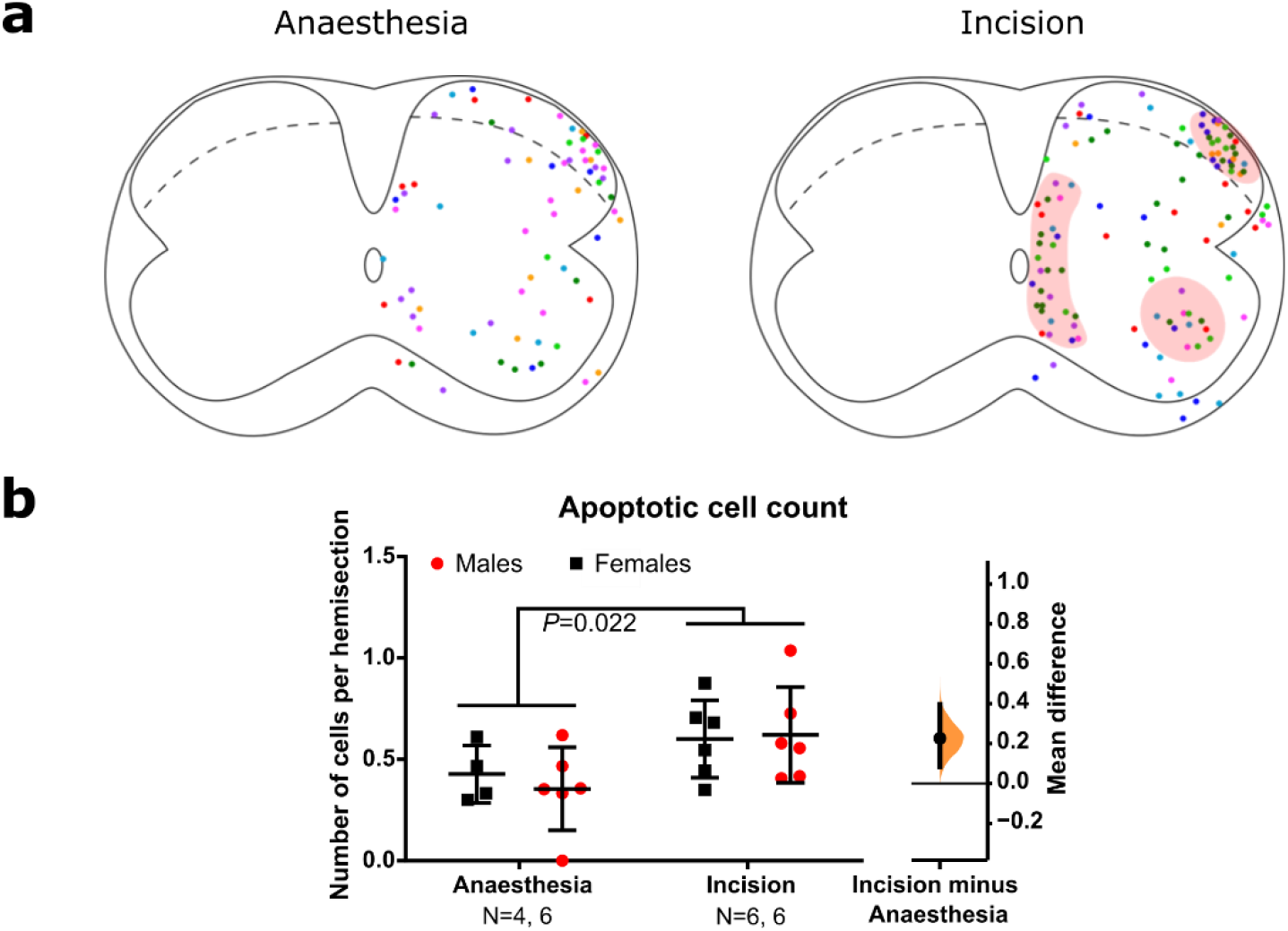
Neonatal incision increases apoptosis in distinct regions within the lumbar spinal cord. **a**. Superimposition of (caspase-3) apoptotic cell locations in the ipsilateral hemisection of 8 control and 8 incision animals (each colour represents one animal, 4 males and 4 females per group). Red shading highlights areas with particularly high apoptotic density. b. Neonatal incision significantly increased the number of apoptotic cells within the ipsilateral lumbar spinal cord. Treatment mean difference 0.23 [95.00% CI 0.085, 0.40]. N-numbers and P-value from two-way ANOVA are indicated in the figure.

### Synaptic engulfment following adult incision and and the influence of prior neonatal incision

Following the characterisation of how neonatal incision affects microglial engulfment of spinal dorsal horn synapses, we investigated microglial engulfment of synapses in the dorsal horn following hindpaw incision in adult animals and the influence of prior neonatal incision (Fig. 1b).

### Neonatal incision decreases VGAT density in adulthood

First, we examined if neonatal incision had any long-term effects on synapse and microglia density in adulthood. For inhibitory presynaptic terminals, neonatal incision decreased VGAT volume (P = 0.0006) (Fig. 4 a, c), indicating a decrease in inhibitory synaptic density, likely a direct consequence of the increased engulfment of inhibitory synapses immediately following neonatal incision. By contrast, excitatory VGLUT2 terminal volume was not altered (P = 0.81).

Despite the neuronal changes, total microglial and lysosomal volume was not altered by neonatal incision in IN(+,−) vs IN(−,−) animals (Fig. 5a), suggesting that neonatal-incision induced increase in microglia density and phagocytic activity is transient.

**Figure 4.**
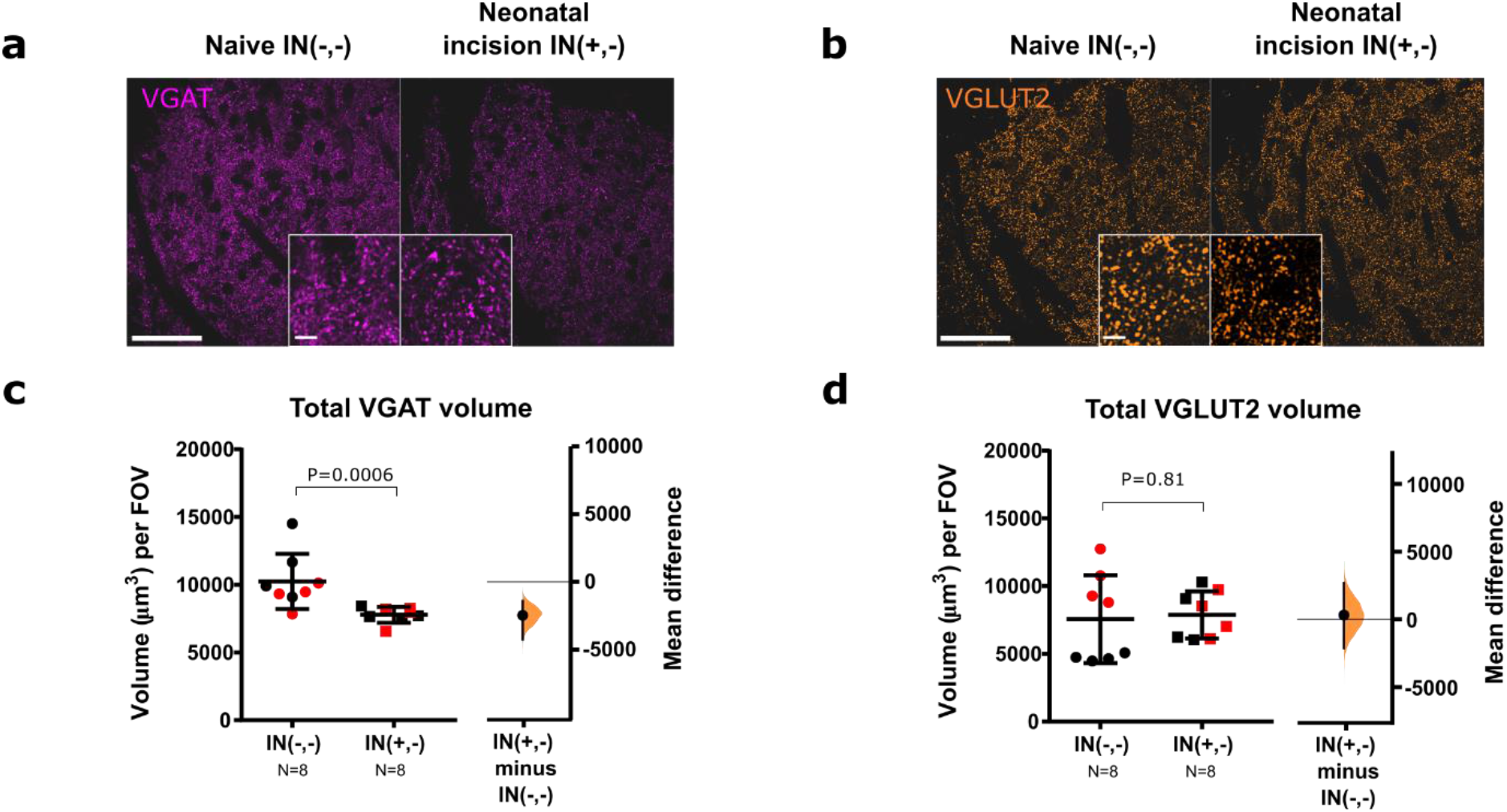
Neonatal incision alters VGAT density long-term. **a., b**. Representative confocal images of VGAT (magenta) and VGLUT2 (orange) staining (scale bar = 50 μm) with insert box showing zoom-in (scale bar = 5 μm). **c**. Total VGAT volume was decreased in adults with neonatal incision vs naïve animals (P= 0.0006, mean difference: −2455.66 [95% CI −4226.98, −1410.13]. **d**. Total VGLUT2 volume was not altered by neonatal incision (P=0.81, mean difference: 317.0 [95% CI −2101.74, 2641.46]). N-numbers and P-values are indicated in Figures. Black and red data points indicate females and males. Field of view (FOV) = 192.79 × 192.79 × 50 μm.

**Figure 5.**
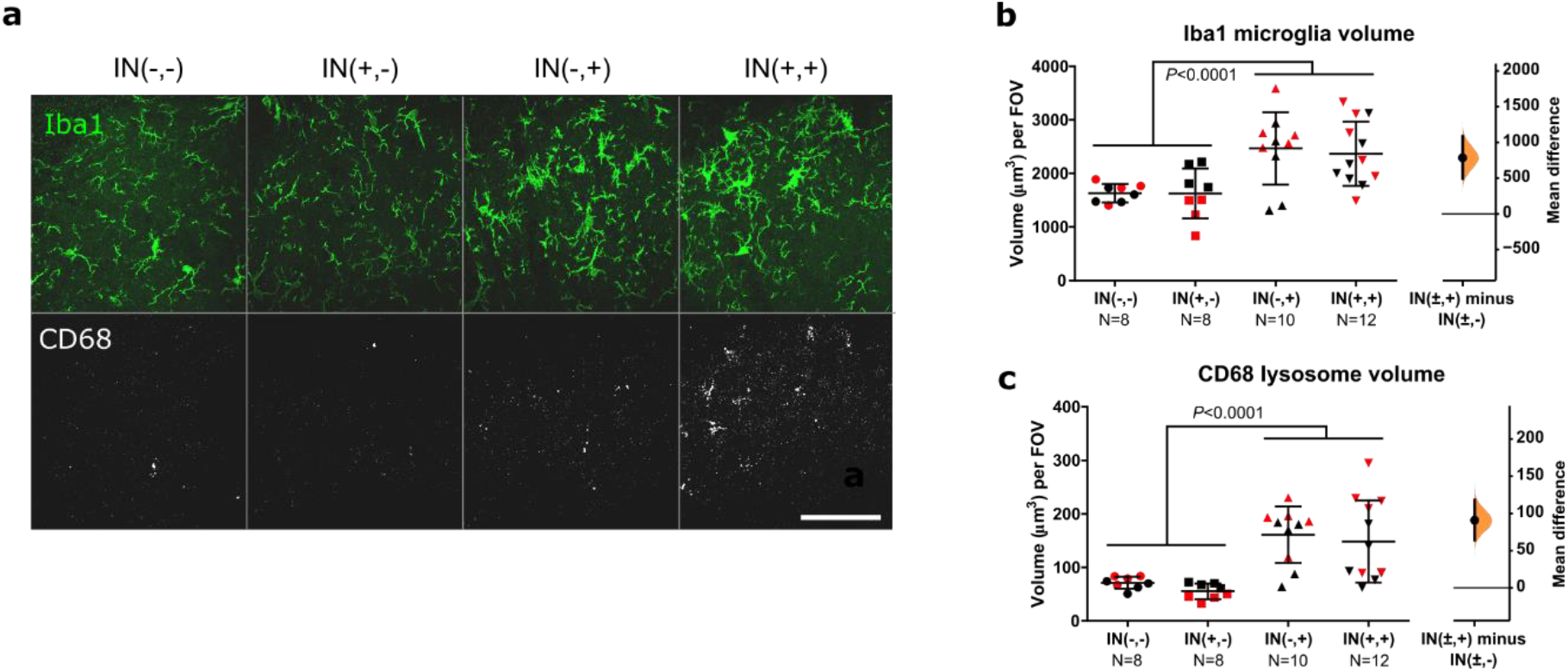
Adult incision increases microglia (Iba1) and lysosomal (CD68) volume regardless of neonatal incision. **a**. Maximum projection of representative confocal images showing microglia (Iba1, green), lysosomes (CD68, grey) for all conditions. IN(−,−): naive, IN(+,−): neonatal incision only, IN(−,+): adult incision only, IN(+,+): neonatal and adult incision. Scale bar = 50μm, Field of view (FOV) = 192.79 × 192.79 × 5 μm. **b**. Microglial Iba1 volume increases following adult incision, mean difference between animals with and without adult incision: 784.22 [95.00% CI 495.94, 1083.49]. **c**. Lysosomal CD68 volume increases following adult incision, mean difference between animals with and without adult incision: 90.83 [95.00% CI 64.17, 118.10]. N-numbers and P-values for two-way ANOVA are indicated in figures. Black and red data points indicate females and males.

### Neonatal incision decreases microglial VGLUT2 engulfment in adulthood

Next, we asked if there is ongoing microglial engulfment of VGAT and VGLUT2 in adult animals and whether this is influenced by prior neonatal incision. VGAT engulfment was not altered (IN(+,−) vs. IN(−,−), P = 0.76) (Fig. 6 a, b), while VGLUT2 engulfment was decreased in animals with prior neonatal incision compared to naïve adults (IN(+,−) vs. IN(−,−), P = 0.041) (Fig. 6 c, d). This suggests that neonatal incision continued to influence how microglia interacted with VGLUT2 synapses in adulthood, although general microglial capacity for engulfment was not changed.

**Figure 6.**
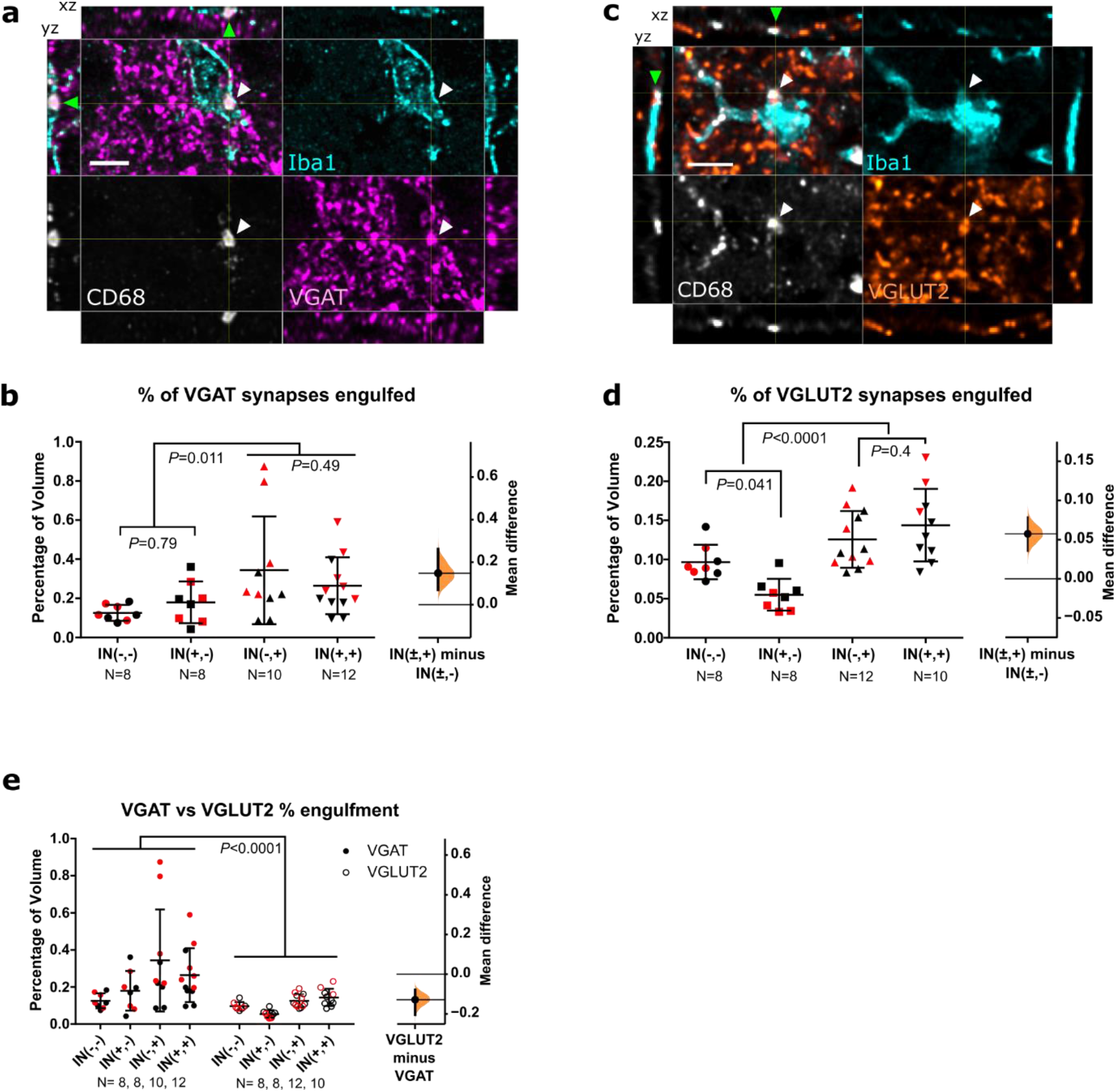
Neonatal and adult incisions differentially affect microglial engulfment of VGAT and VGLUT2 synapses. **a., c**. Representative confocal images from the superficial medial lumbar dorsal horn showing (a) inhibitory (VGAT, magenta) and (c) excitatory (VGLUT2, orange) presynaptic terminals with microglia (Iba1, cyan) and lysosomes (CD68, grey). White and green arrowheads point out overlap of Iba1, CD68, and VGAT in the xy view and side panels respectively, indicating engulfment. Cross-hairs show position of the xz and yz side-view panels. Scale bar = 5μm. **b**. Microglial engulfment of VGAT presynaptic terminals increases following adult incision, mean difference between animals with and without adult incision: 0.147 [95.00% CI 0.069, 0.261]. **d**. Neonatal incision decreases baseline microglial engulfment of VGLUT2 in adulthood, while adult incision increases VGLUT2 engulfment irrespective of neonatal incision. Mean differences: neonatal incision only vs naive (light grey, IN(+,−) minus IN(−,−)): −0.042 [95.00% CI −0.062, −0.022]; adult & neonatal incision vs adult incision only (dark grey, IN(+,+) minus IN(−,+)): 0.018 [95.00% CI −0.014, 0.054]. **e**. Microglial preferentially engulf inhibitory VGAT presynaptic terminals (comparing data from b & d). Mean difference −0.13 [95.00% CI −0.20, −0.08]. N-numbers and P-values from two-way ANOVA (with Sidak posthoc) are indicated in figures. Black and red data points indicate females and males.

### Neonatal incision does not influence microglial synaptic engulfment following adult re-incision

Next, we examined the effect of adult incision and any interaction with prior neonatal incision on microglia and microglial engulfment of inhibitory (VGAT) and excitatory (VGLUT2) synapses. As with neonates, adult incision (IN(±,+)) increased Iba1 microglial volume compared to animals without adult incision (IN(±,−)) (adult incision effect F (1, 34) = 19.88, P < 0.0001) (Fig. 5a, b). This was irrespective of whether the animals had prior neonatal injury (neonatal incision effect F (1, 34) = 0.088, P = 0.79; interaction F(1,34) = 0.067, P = 0.80). The same effect was observed for CD68 lysosomal volume within microglia (Fig. 5a, c). Adult incision increased CD68 lysosomal volume compared to animals without adult incision regardless of neonatal incision history, suggesting that microglial phagocytosis following an adult incision was generally increased, but not influenced by neonatal incision (adult incision effect F (1, 34) = 28.48, P < 0.0001; neonatal incision effect F (1, 34) = 0.70, P = 0.41; interaction F (1, 34) = 0.01, P = 0.92).

We then examined the proportion of VGAT and VGLUT2 synapses engulfed by microglia. Adult incision increased the percentage of engulfed synapses for both VGAT and VGLUT2 (VGAT: F (1, 34) = 7.149, P = 0.011; VGLUT2: F (1, 34) = 27.30, P < 0.0001 comparing IN(±, +) vs IN(±, −)). For VGAT, this was irrespective of whether animals had prior neonatal incision or not. Interestingly, for VGLUT2 there was an interaction (F (1, 34) = 7.018, P = 0.0122), as neonatal incision decreased the percentage of baseline VGLUT2 engulfment in adulthood (P=0.041 IN(−,−) vs IN(−,+)), but microglia in both groups engulfed VGLUT2 synapses to the same extend after adult re-incision (P=0.40 IN(−,+) vs IN(+,+)). As adults with or without neonatal incision both responded similarly to an acute adult incision (IN(+,+) vs IN(−,+)) in terms of changes in microglial volume, CD68 volume, and the percentage of engulfed VGAT and VGLUT2, this suggests that neonatal incision did not train microglia cells in their phagocytic response.

### Adult incision predominantly increases microglial engulfment of inhibitory synapses

We then investigated whether microglial engulfment had any synapse specificity by comparing total percentage of engulfed VGAT and VGLUT2 synapses. The percentage of VGAT engulfment was considerably higher than VGLUT2 engulfment (Two-way ANOVA, F(1; 68) = 18:10, P < 0.0001) (Fig. 6e), and therefore microglial preferentially targeted inhibitory synapses. In contrast to neonatal incision, adult incision had no effect on microglial A-fibre engulfment and was not affected by prior neonatal incision (one-way ANOVA (F(2,27) = 0.92, P = 0.41) (Fig. S1).

## Discussion

It is well known that peripheral incision in neonates and adults can alter dorsal horn processing acutely and long-term, but how this is achieved is unclear. Here we show that microglia contribute to this process through the engulfment of presynaptic terminals following incision.

Both adult and neonatal microglia show a preference for the engulfment of local inhibitory VGAT terminals, with neonatal microglia additionally engulfing A-fibre terminals following incision. Neonatal incision reduced inhibitory VGAT density long-term while VGLUT2 density was not affected. However, neonatal incision decreased microglial phagocytosis of VGLUT2 terminals in adulthood, suggesting that although their density was not altered, there might be a reduction in their turn-over rate. Neonatal incision did not affect microglial phagocytosis of synapses in response to re-incision in adulthood, which suggests that microglial phagocytosis did not become trained. Overall, the results support our first hypothesis that altered microglial engulfment of neurites following incision leads to long-term structural changes which could underlie the behavioural changes observed in clinal and animal models following early postnatal injury.

### Acute consequences of incision in neonates

In neonates, there was no statistical interaction between sex and incision. This suggests that both males and females responded similarly to incision. However, females tended to have higher microglia volume, and subsequently higher lysosome volume and synaptic engulfment in all conditions. In contrast to our finding, Schwarz et al., (2012) reported that males have higher microglial density and more reactive phenotype in several brain regions at baseline at P4, which suggests that brain and spinal cord microglia are differentially regulated.

It has been shown that neonatal-incision induced hyperalgesia in adulthood is mediated by microglial p38 activation in males but not in females despite both males and females showing increased microglial proliferation following incision (Moriarty et al., 2019). This suggests that some aspects of microglial response towards incision – including phagocytosis – are shared between males and females while other aspects differ.

In contrast to adult incisions, neonatal incision only caused a modest increase in Iba1 volume and no significant changes in lysosome volume despite enhanced engulfment of inhibitory synapses and A-fibres following incision. This is likely due to the lower microglia density and reactivity in neonates consistent with previous results (Moss et al., 2007).

Neonatal incision selectively increased microglial engulfment of inhibitory VGAT synapses but not excitatory VGLUT2 synapses, both of which are primarily expressed by local interneurons (Punnakkal et al., 2014). Previous reports of microglial engulfment in the postnatal brain and spinal cord have primarily focused on excitatory synapses (Paolicelli et al., 2011; Schafer et al., 2012b; Vainchtein et al., 2018; Vukojicic et al., 2019), while report of inhibitory synapses has been sparse (Favuzzi et al., 2021). Our finding adds to the accumulating evidence that inhibitory synapses are also engulfed and modulated by microglia during development. Interestingly, Salter et al. (2020) have shown that synapse engulfment in the neonatal hippocampus was specific to VGLUT2 but not VGLUT1 or VGAT synapses. Together, this suggests that microglial engulfment is not only specific to synapse identity but also varies between CNS regions.

A-fibre terminal distribution and inhibitory circuits mature and undergo plastic changes over the first postnatal week (Koch et al., 2012; Koch & Fitzgerald, 2013), with microglia actively engulfing A-fibres as part of normal development (Xu et al., 2021). This might render them more susceptible to external perturbations and could explain why A-fibre and VGAT presynaptic terminals are engulfed following peripheral injury, but VGLUT2 presynaptic terminals are not.

An over-pruning of A-fibres following incision will likely alter the pattern of tactile input to the dorsal horn and affect behavioural sensitivity long-term. This is supported by our recent finding that microglial dysfunction in neonates lead to structural, functional and behavioural changes in adults (Xu et al., 2021). Alternatively, the engulfment of A-fibres could be a homeostatic response to A-fibre sprouting, which has been observed in neonatal animals following nerve transection and hindpaw inflammation (Fitzgerald, 1985; Fitzgerald et al., 1990; Ruda et al., 2000) and is supported by electrophysiology studies showing that neonatal incision increased peripheral afferent drive and monosynaptic A-fibre inputs onto lamina I projection neurons in adulthood (Li et al., 2015).

VGAT synapses in the dorsal horn could belong to local interneurons or descending input from the RVM (cortico-spinal inputs do not arrive in the grey matter of the lumbar cord until P9 (Donatelle, 1977)). While the net effect of GABA and glycine on the neonatal spinal cord is excitatory (Hathway et al., 2006; Koch et al., 2012), the depolarising effect is presumed to be mainly important for the maturation of the spinal cord rather than providing an excitatory drive (Baccei & Fitzgerald, 2004). Therefore, excessive pruning of VGAT synapses could impair the normal maturation of local inhibitory circuits.

As microglia are known to induce apoptosis and clear apoptotic cell debris, we investigated whether apoptosis might be altered in the medial dorsal horn region innervated by the hindpaw and where microglial proliferation is observed following hindpaw incision.

Consistent with previous reports (Moriarty et al., 2019), apoptotic cell count in the ipsilateral spinal cord increased following neonatal incision, but there was no accumulation of apoptotic cells in the medial dorsal horn. Instead, apoptotic cells aggregate in the superficial lateral dorsal horn, the ventral horn, and along the central canal. This suggests that the incision-induced apoptosis is likely not due to direct microglial activity. It is possible that hindpaw incision interferes with microglial removal of apoptotic cells, and reveals the normal ongoing rate of apoptosis, which under normal circumstances would have been rapidly removed by microglia. This is consistent with the distribution of apoptotic profiles in naïve animals being similar but reduced in number.

#### Long-term consequences of neonatal incision in adulthood

Consistent with the increase of microglial VGAT engulfment following neonatal incision, we observed a reduction of VGAT density in adulthood. Impaired inhibitory input is a common feature of various pain states (Baba et al., 2003; Drew et al., 2004; Moore et al., 2002; Sivilotti & Woolf, 1994; Zimmer et al., 2004). Therefore, a reduction of inhibitory VGAT terminals in the superficial dorsal horn could contribute to hyperalgesia upon re-incision in adulthood. This supports our first hypothesis that neonatal incision alters microglial phagocytosis, leading to long-term structural changes that could underly behavioural deficits in adulthood.

In addition, neonatal incision altered microglial interaction with VGLUT2 terminals long-term, leading to decreased VGLUT2 engulfment in adulthood. As total VGLUT2 density was not altered, the reduced phagocytosis might indicate a slower turnover rate for VGLUT2 synapses. It is known that neonatal incision decreases baseline sensitivity in adulthood which is mediated by a reduction of primary afferent drive and increase of inhibitory influence from the brainstem (Grunau et al., 1994; Schmelzle-Lubiecki et al., 2007; Walker et al., 2016), and it is possible that altered microglia-VGLUT2 interaction either contribute to this phenomenon, or is altered in compensation to the reduced sensitivity.

Despite the difference in microglial engulfment for VGLUT2 at baseline, neonatal incision did not affect microglial engulfment of VGLUT2 or VGAT synapses after adult incision, i.e. animals were able to mount a normal phagocytic response regardless of neonatal incision. Therefore, neonatal incision did not train microglial phagocytosis. However, phagocytosis is not the only microglial interaction with neurons and synapses, and other aspects of microglial physiology (e.g. cytokine release) were trained or altered that contribute to the behavioural hyperalgesia in adulthood.

#### Acute consequences of incision in adults

In adults, we also saw a preferential engulfment of inhibitory VGAT terminals following incision, while VGLUT2 terminals were engulfed to a lesser degree and A-fibre engulfment was not altered. Previous studies on microglial engulfment in adulthood have either focused on excitatory synapses or used pan-synaptic markers that do not distinguish between excitatory and inhibitory synapses (Hong et al., 2016; Shi et al., 2015; Stevens et al., 2007; Wang et al., 2020), potentially missing synapse-specific interactions as observed here. The preferential engulfment of VGAT synapses might be due to differences in neuronal activities between VGAT and VGLUT2 synapses. Inhibitory neurons are known to have tonic activity (Yasaka et al., 2010). Given that microglia can sense neuronal activity and tend to contact highly active synapses more often, this might explain why a higher percentage of VGAT synapses are engulfed (Eyo et al., 2016; Kato et al., 2016; Wake et al., 2009). Inhibitory synapses are proposed to be modulators of network activity, as they can both inhibit fast transmission via synaptic release and set the inhibitory tone via extra-synaptic release sites (Farrant & Nusser, 2005; Lee & Maguire, 2014). Therefore, the modulation of inhibitory synapses in adults might directly affect network activity in the dorsal horn. Further, altered balance of excitation vs inhibition is implicated in neuropathic pain in the form of disinhibition (Latremoliere & Woolf, 2009). Therefore, the higher loss of inhibitory VGAT synapses could cause disinhibition within the dorsal horn and contribute to hypersensitivity following incision. This is supported by studies showing that pharmacological inhibition of spinal GABA and glycine signalling or facilitation of glutamate signalling can both result in hypersensitivity and altered sensory processing (Liaw et al., 2005; Yaksh, 1989). In animal with prior neonatal incision, this effect is likely compounded by the lower density of VGAT and could explain the hyperalgesia observed following re-incision in adulthood. In contrast to neonates, A-fibre engulfment was not altered following hindpaw incision in adults. Previous reports have shown that hyperalgesia could only be induced if injury happened within a critical period (1st postnatal week) (Walker et al., 2009), which coincides with the maturation period for A-fibres (Xu et al., 2021). Therefore, it is possible that A-fibres are protected from engulfment once the critical period is closed. This is further supported by a recent study on microglial engulfment of synapses following adult peripheral nerve injury after which VGAT and VGLUT2 synapses were engulfed 20 days post injury, but not afferent VGLUT1 or VGLUT3 synapses (Yousefpour et al., 2023). This suggests that the type of synapses engulfed is conserved following different types of peripheral injury, but the extent and timing of engulfment can differ.

#### Limitations

For presynaptic terminals, it has also been shown that microglia are capable of engulfing parts of it without removing the whole synapse (Weinhard et al., 2018). This suggests that microglia are capable of more subtle modulations of synapses apart from complete removal. It is also not completely clear whether this modulation could be strengthening synapses instead, but there is evidence that repeated microglial contact with spines causes their disappearance and presumably loss of any synapse they were a part of (Tremblay et al., 2010; Wake et al., 2009). Therefore, although evidence does point to microglial removal of synapses, other interactions cannot be excluded. However, differences in the amount of microglial contact, regardless of whether actual engulfment has happened, is in itself an indication that there are differences in microglial behaviour. Therefore, this does not change the conclusion that incision alters microglial modulation of presynaptic terminals.

## Conclusions

Previous research has suggested that microglia could be a promising therapeutic target in pain interventions (Wen et al., 2011). However, microglial function in the spinal cord in relation to pain has mainly been characterised in terms of proliferation, cytokine production, and activation of the p38 and P2X4 pathway (Taves et al., 2013; Wen et al., 2011). Here we show that microglial phagocytosis of synapses is part of the physiological response to incision injury, and that they preferentially engulf inhibitory synapses in adulthood, which could provide another target for pain intervention. As adult and neonatal dorsal horn microglia react differently to hindpaw incision, any strategies targeting microglia cells will require different approaches in neonates and adults, while taking their developmental functions in neonatal spinal cord into consideration.

## Acknowledgements

The authors thank the Biological Services Unit at UCL for support in animal maintenance and Stuart Martin for genotyping.

## Author Contributions

YX and WJ executed experiments, YX and DM wrote Fiji macros for analysis, YX and SB designed experiments and wrote the manuscript.

---

**Figure S1.**
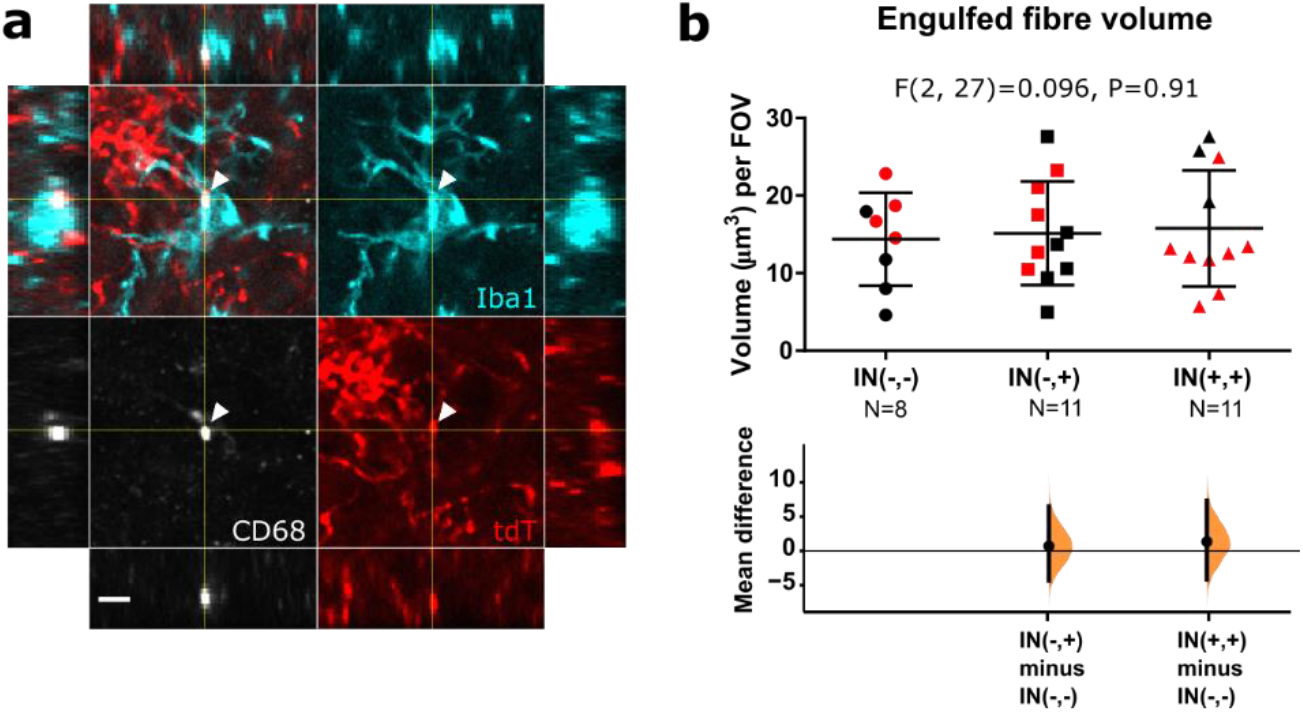
Adult incision did not alter A-fibre engulfment. **a**. Representative confocal image showing microglia (Iba1, cyan), lysosomes (CD68, grey), and A-fibres (tdT, red). White arrowheads point out overlap of Iba1, CD68, and tdT, indicating engulfment. Cross-hairs show position of the xz and yz side-view panels. Scale bar = 5μm. **b**. Engulfment of A-fibres is not altered following adult incision regardless of neonatal incision. Mean difference between adult incision only and naive animals (IN(−,+) minus IN(−,−)): 0.75 [95.00% CI −4.26, 6.52], mean difference between adult incision with neonatal incision and naive animals IN(+,+) minus IN(−,−)): 1.39 [95.00% CI −4.09, 7.36]. FOV = 192.74 × 192.74 × 50 μm. N-numbers and P-values are indicated in Figures. Black and red data points indicate females and males.

